# Inkjet based 3D Printing of bespoke medical devices that resist bacterial biofilm formation

**DOI:** 10.1101/2020.06.30.180596

**Authors:** Yinfeng He, Belen Begines, Jeni Luckett, Jean-Frédéric Dubern, Andrew L. Hook, Elisabetta Prina, Felicity R.A.J. Rose, Christopher J. Tuck, Richard J.M. Hague, Derek J. Irvine, Paul Williams, Morgan R. Alexander, Ricky D. Wildman

## Abstract

We demonstrate the formulation of advanced functional 3D printing inks that prevent the formation of bacterial biofilms *in vivo*. Starting from polymer libraries, we show that a biofilm resistant object can be 3D printed with the potential for shape and cell instructive function to be selected independently. When tested *in vivo*, the candidate materials not only resisted bacterial attachment but drove the recruitment of host defences in order to clear infection. To exemplify our approach, we manufacture a finger prosthetic and demonstrate that it resists biofilm formation – a cell instructive function that can prevent the development of infection during surgical implantation. More widely, cell instructive behaviours can be ‘dialled up’ from available libraries and may include in the future such diverse functions as the modulation of immune response and the direction of stem cell fate.

## Introduction

Medicine is moving towards meeting the needs of individual patients through patient stratification and personalisation. This can arise in many forms e.g. for implanted medical devices, through tailoring of shape, of material and of function, each tuned to the specific needs of the patient. One method to achieve this is through ‘additive manufacturing’ (AM) or ‘3D Printing (3DP)’ ^[1–3]^ but it requires the identification and scale-up of 3D printable materials or formulations that have the required functional performance. We propose a strategy that allows us to formulate materials with cell instructive properties that adds an additional lever of personalisation to current 3D printed devices. We exemplify this strategy by testing the efficacy of our formulations *in vitro, in vivo* and by manufacturing a bespoke finger joint prosthetic that demonstrates resistance to bacterial biofilm formation. This choice of cell-instructive function is inspired by the need to prevent medical device associated infections ^[4–7]^. Studies have shown that 1 to 5% of implanted prostheses become colonized ^[8, 9]^ with biofilms formed by pathogens such as *Pseudomonas aeruginosa* and *Staphylococcus aureus* resulting in poor clinical outcomes ^[10]^. Such biofilms are refractory to both antibiotic therapy and to clearance by host immune defence mechanisms leading to chronic infections and device failure. Thus, the use of personalised prosthetics brings with it a need to prevent microbial infection. Whilst our demonstrator focuses on infection prevention, our manufacturing strategy is agnostic with respect to the cell instructive functionality and material libraries.

The chosen demonstrator device differs in approach to other anti-microbial devices, such as those blending antibiotics into the material or through surface modification ^[11–18]^. These approaches have multiple issues including coating delamination and cracking in the aggressive implant service environment ^[16]^, localized cytotoxicity from anti-microbial coatings ^[16]^, active compound depletion^[17,18]^, and most crucially, potential selection for anti-microbial resistance resulting from the selective pressures that antimicrobial killing strategies impose^[19]^. Here we exploit acrylate monomers, previously identified to prevent bacterial biofilm formation^[20–22]^ and use these as the basis for inks for inkjet based 3D printing (IJ3DP), an AM technique whose strengths come into play when there is a need for industrial scalability, high resolution, and multi-material manufacture. A biofilm prevention strategy reduces the evolutionary pressures that drive the development of antimicrobial resistance. The biofilm surface coverage on polymer samples was reduced by 99% compared with those on silicone rubber for diverse multi-antibiotic resistant pathogens including *P. aeruginosa*, *S. aureus Escherichia coli, Klebsiella pneumoniae, Enterococcus faecalis and Proteus mirabilis* ^[22, 23]^. Here, we present a novel approach that shows how the candidate materials identified from such a screen can be adapted to create formulations suitable for IJ3DP and demonstrate that these formulations are effective *in vitro* and *in vivo in a foreign body infection model*. We also ascertained that the mechanical and cell instructive performance of the materials is intimately tied to the degree of polymerization during printing and that there was no evidence of bacteria being killed through contact with the material or the leaching of any toxic residuals.

This approach, illustrated graphically in Fig. 1, takes materials from a previously published polymer library based on a screening for resistance to bacterial attachment and biofilm formation. We assessed candidates from the library for their capacity for consistent and reliable deposition from an ink jet print head. Formulations showing promise were used to create specimens that were characterised through a series of tests that examined: 1. level of uncured acrylates, 2. mechanical properties, 3. bacterial biofilm formation and 4. cytotoxicity towards bacterial cells. Since a potential application is to produce devices that may be used in a clinical context, we also investigated their mammalian cell cytotoxicity (following ISO 10993 guidelines) as well as *in vivo* biofilm inhibition performance tested in a mouse infection model. Thus, the formulations with the most promising performance were identified, used to produce concept devices by IJ3DP and challenged to understand their efficacy following manufacturing.

**Fig. 1:**
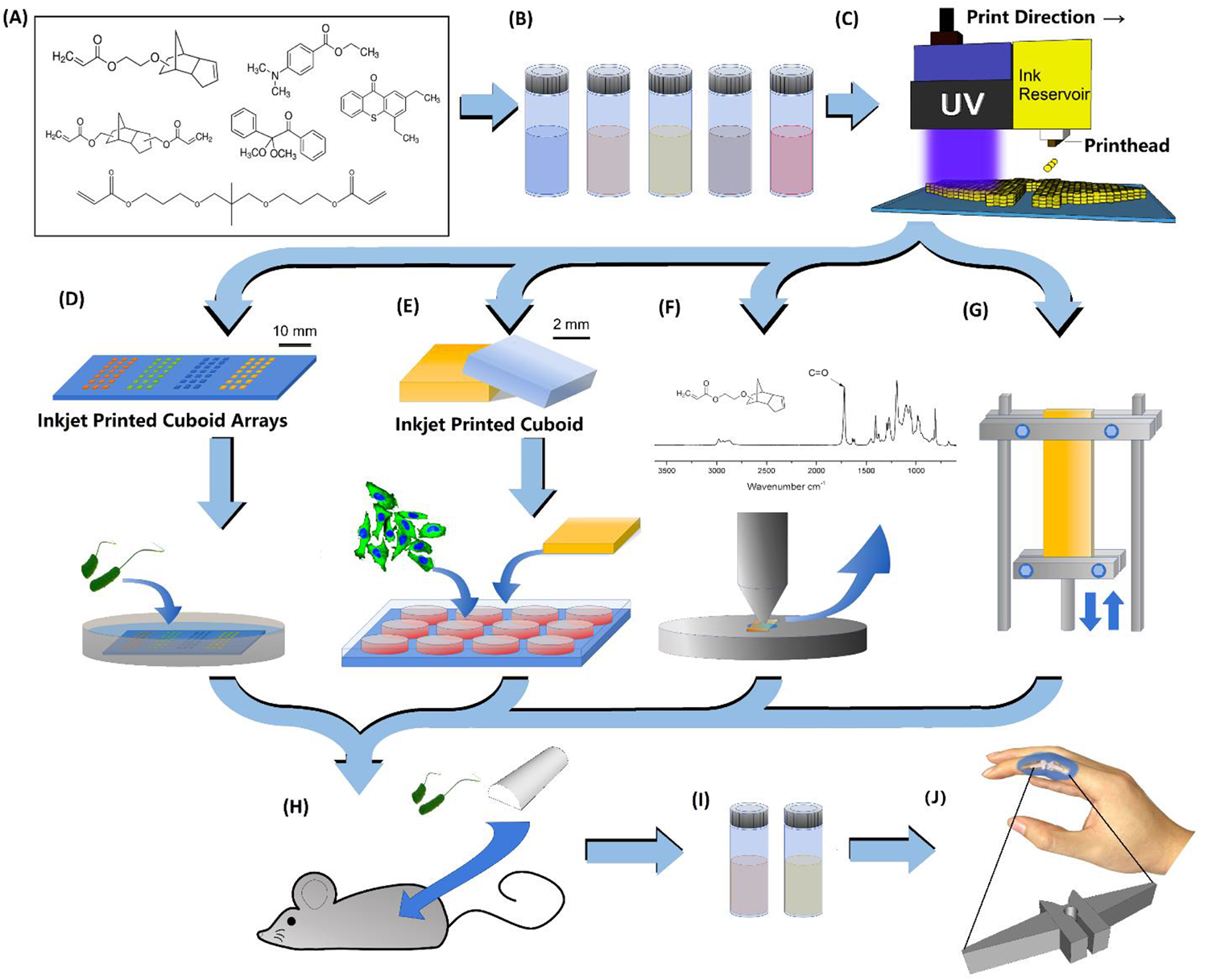
Schematic for developing optimized formulations for IJ3DP. **A-B)** Selected monomer candidates for the formulation development and optimization. **C)** A Fujifilm Dimatix DMP-2830 3D printer was used to print samples. The system in this case was equipped with a cartridge ejecting 10 pL drop volumes, utilising up to 16 nozzles. **D)** On slide arrays of cuboids were created by IJ3DP for preliminary microbiology biofilm assays using P. aeruginosa. **E)** Cytotoxicity and cell attachment biocompatibility tests on the printed samples were carried out using mouse embryonic fibroblast 3T3 fibroblasts to assess biocompatibility of the printed device; **F)** Attenuation Total Reflectance Infrared Spectroscopy (ATR-IR) was used to quantify the levels of residual acrylate in the specimens made from different ink formulations; **G)** Mechanical tests were performed by Dynamic Mechanical Analysis (DMA) in tension mode at room temperature; **H)** Formulations resulting in acceptable properties were tested in vivo to ensure the cell instructive retained in a more complex environment; **I-J)** The finalized ink formulations were used to print concept devices.

## Results

Using published lists of materials resistant to bacterial attachment we selected suitable candidates first using printability as a guide^[24]^, based on physical characteristics such as viscosity and surface tension, screening out those materials that were outside the range commonly accepted as ‘printable’ for inkjet. A trial printing of the remaining materials was then conducted to determine the reliability of printing and whether the materials would cure on our printing configuration. From this we selected two materials that showed the greatest promise for successful printing and proceeded to optimise the formulations ready for scale up^[25–28]^ (Supplementary Table S1): tricyclo[5.2.1.02,6]decanedimethanol diacrylate (TCDMDA) and ethylene glycol dicyclopentenyl ether acrylate (EGDPEA). Sixteen formulations were then investigated where the photoinitiator and the candidate monomers were combined covering a breadth of utility in different environments and potential reaction speeds (Supplementary Table S2) as both could influence the product performance^[22, 29–31]^. Both Norrish type I (nitrogen environment) and Norrish type II (air environment) initiators were evaluated with respect to compatibility of the formulations when processing in different environments. A series of tests on each of our formulations were conducted to understand the performance of our 3D printed constructs *in vitro* and *in vivo*.

### Bacterial Biofilm Formation on Polymer Cuboid Arrays

To determine whether printed samples retained their desired biofilm resistance, samples were printed using a laboratory-based inkjet printer. For each formulation, the printed samples consisted of a series of 24 cuboid arrays (2000 x 2000 x 100 μm^3^ each) (Fig. 2A) on polystyrene slides. *P. aeruginosa* biofilm surface coverage on the polymers was quantified after culturing for 72 h. All the inkjet printed and cured poly-EGDPEA and poly-TCDMDA surfaces showed low biofilm surface coverage when compared with the silicone rubber control (Appleton Woods medical grade tubing). In previous work, poly-EGDPEA and poly-TCDMDA samples were prepared by monomer casting then curing and showed significant biofilm resistance, resulting in low *P. aeruginosa* biofilm coverages of 4.0% ± 1.5% and 2.3% ± 1.3% respectively ^[21]^. The poly-EGDPEA and poly-TCDMDA made from inkjet printable formulations in this study showed a bacterial coverage of 3.0 ± 0.9% and 3.3% ± 1.2% respectively compared with >30% for silicone (Fig. 2A), suggesting the performance of the IJ3DP samples was not statistically significantly different from those that had been cast and photopolymerized (*p* = 0.26 and *p* = 0.27) and indicates that both materials retained their ability to prevent bacterial biofilm formation after IJ3DP.

**Fig. 2:**
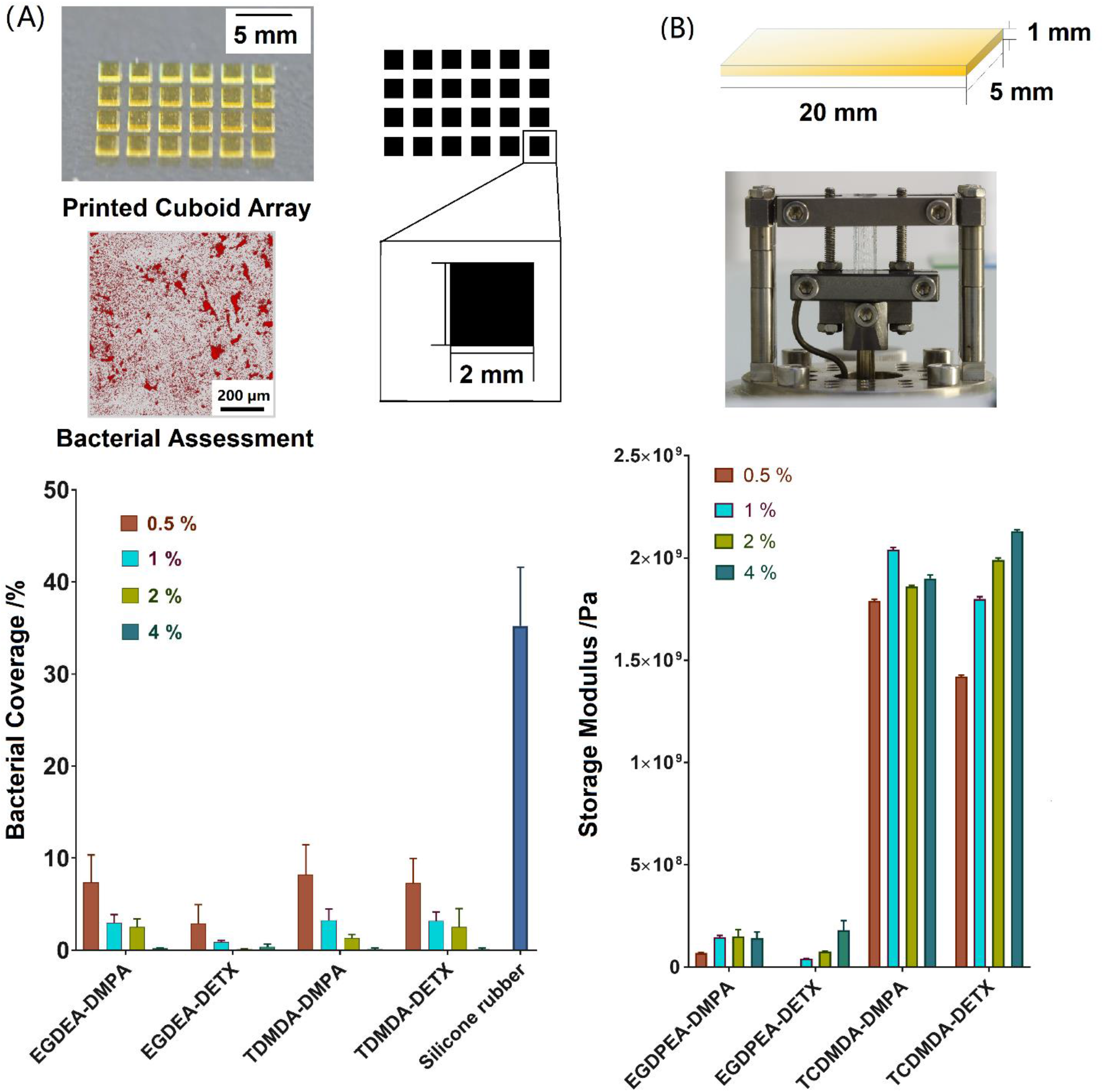
Comparison of the mechanical and bacterial biofilm inhibition performance between IJ3DP printed specimens using selected ink formulations: **A)** An array of 24 cuboids (2 mm x 2 mm x 0.1 mm; w x l x h) was printed onto polystyrene slides and bacterial biofilm formation compared with a silicone control; the samples were imaged after incubation with *P. aeruginosa* using confocal microscopy and biofilm surface coverage was assessed over 640 x 640 μm and presented as surface coverage (%) over the whole assessment window (Mean ± Standard Deviation, n = 24); **B)** The storage modulus of specimens made from all the formulations were measured by dynamic mechanical analysis using strip samples printed (5 mm x 20 mm x 1mm (w x l x h)) (Mean ± Standard Deviation, n = 5).

Specimens formed using formulations containing higher concentrations of photoinitiator were found to result in lower bacterial biofilm coverage. For example, the bacterial surface coverage on the poly-TCDMDA decreased from 8.2% ± 3.2% to 0.2% ± 0.1% when going from 0.5 wt% to 4 wt% DMPA initiator concentration. This reduction in biofilm coverage is attributed to higher conversion of polymer, suggesting the material’s ability to resist biofilm formation is enhanced as conversion is increased. The alternative explanation that the increased photoinitiator concentration correlates with cytotoxicity of the material towards bacteria was ruled out using bacterial growth assays that are presented in Fig. 6A and Supplementary Fig. S4.

**Fig. 3:**
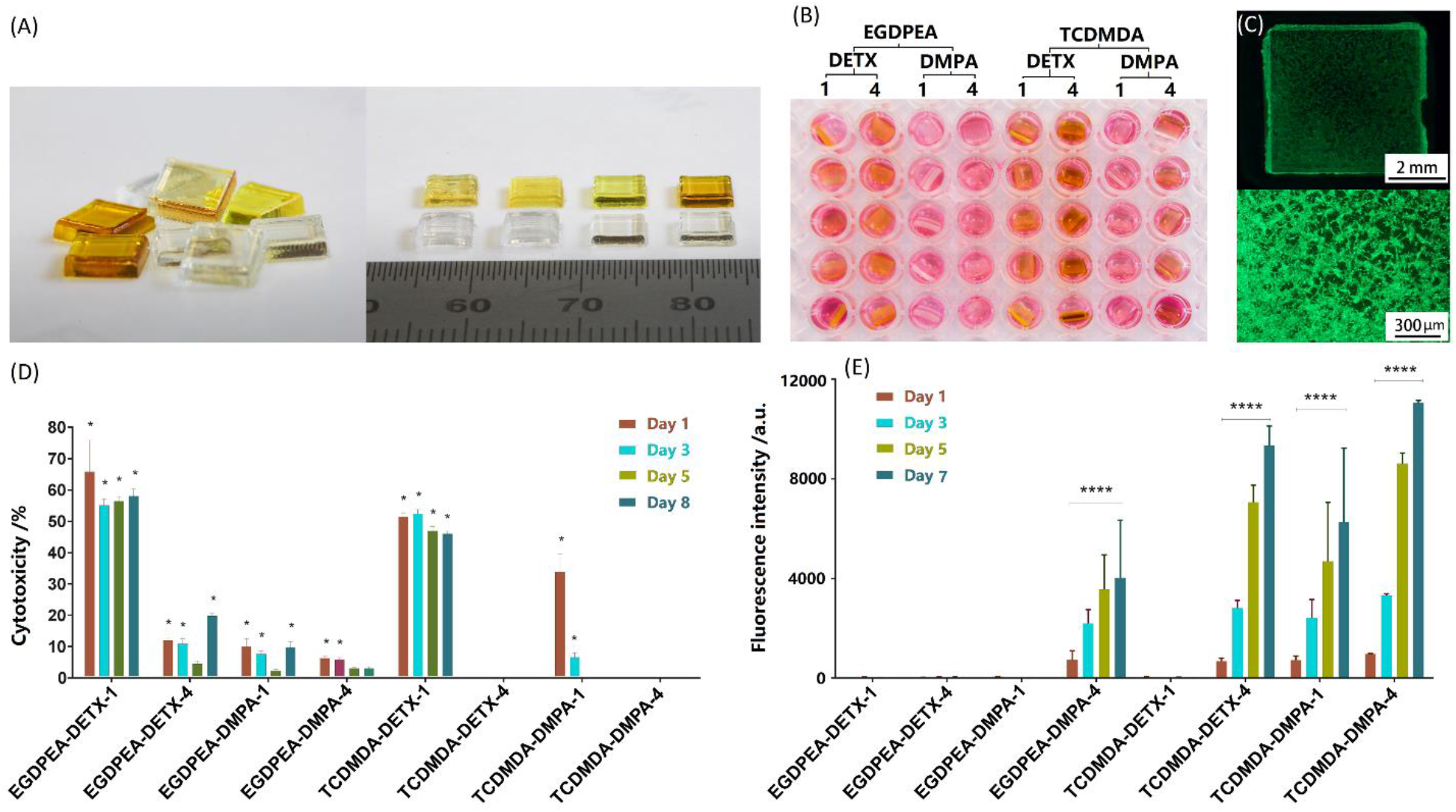
Mammalian 3T3 cell cytotoxicity and attachment tests on printed devices. **A)** Cuboid tablets were printed with the ink formulations in Table S1. **B)** Samples were immersed in culture medium over time (1, 3, 5, and 8 days), and samples transferred to plates containing adherent 3T3 cells. **C)** An example of the Live/Dead^®^ cell viability assay, which in this case was performed on a poly-TCDMDA-DMPA-4 sample illustrating viable cells and proliferation. **D)** Comparison of cytotoxicity (%) for the printed cuboid tablets on different days performed using the LDH assay. Mean ± Standard Deviation with n = 5. **E)** Fluorescence intensity of 3T3 cells seeded on the samples in different formulations measured using the Presto Blue assay. The cells adhered and proliferated on 4 formulations (EGDPEA-DMPA-4, TCDMDA-DETX-4, TCDMDA-DMPA-1, and TCDMDA-DMPA-4); cells cultured on TCDMDA-DMPA-4 demonstrated the highest cell metabolic activities at day 7. Mean ± Standard Deviation, n = 5 (*p ≤ 0.05).

**Fig 4:**
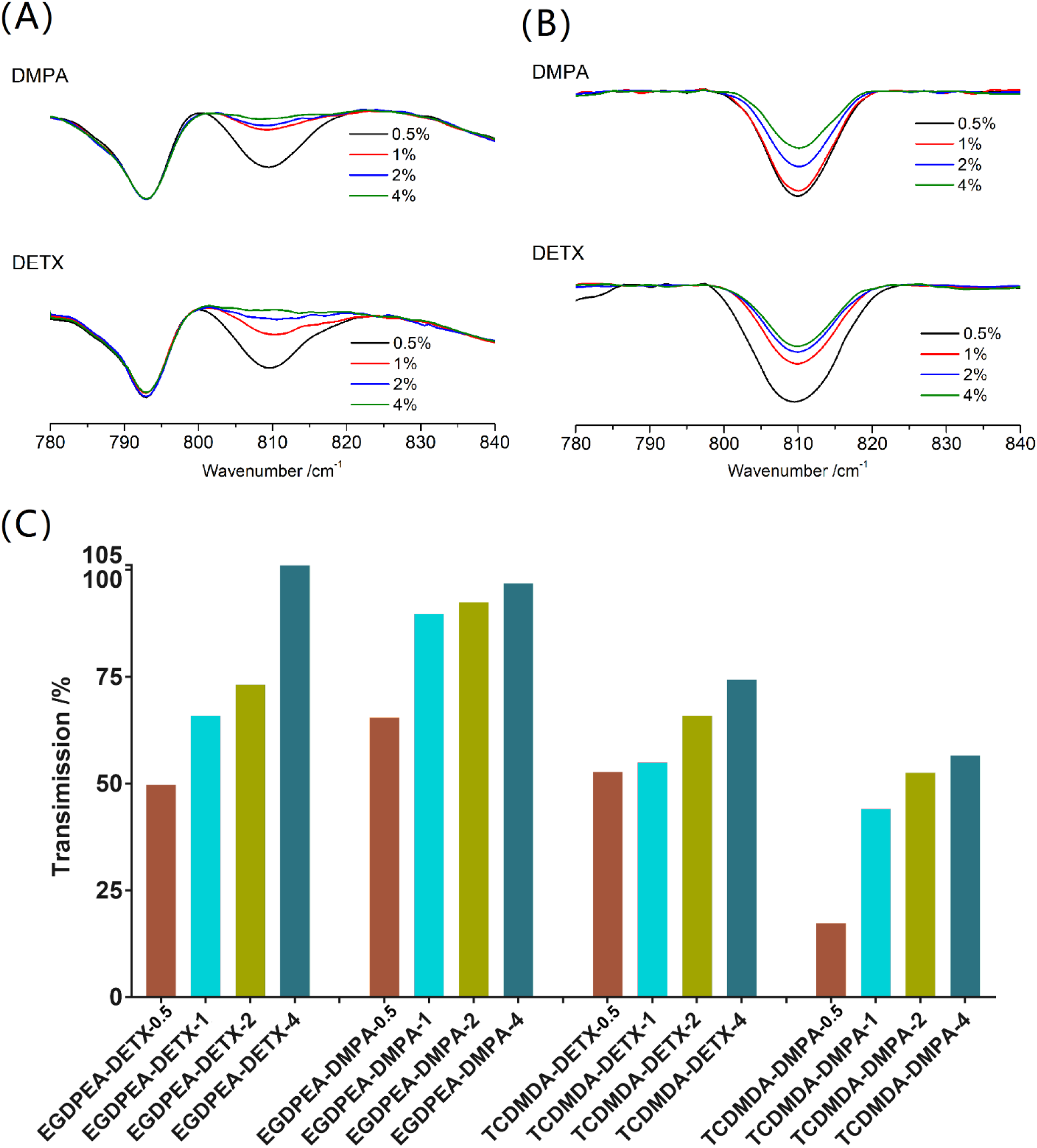
analysis highlighting the trends in residual alkene groups representing the residual monomer and inversely proportional to the level of conversion: The 810 cm^-1^ absorption assigned to the C-H out-of-plane bending vibration of the alkene group displayed normalised to the background intensity. The materials interrogated were **A)** poly-EGDPEA, and **B)** poly-TCDMDA; **C)** Plot of the transmission peak height at 810 cm^-1^.

**Fig.5.**
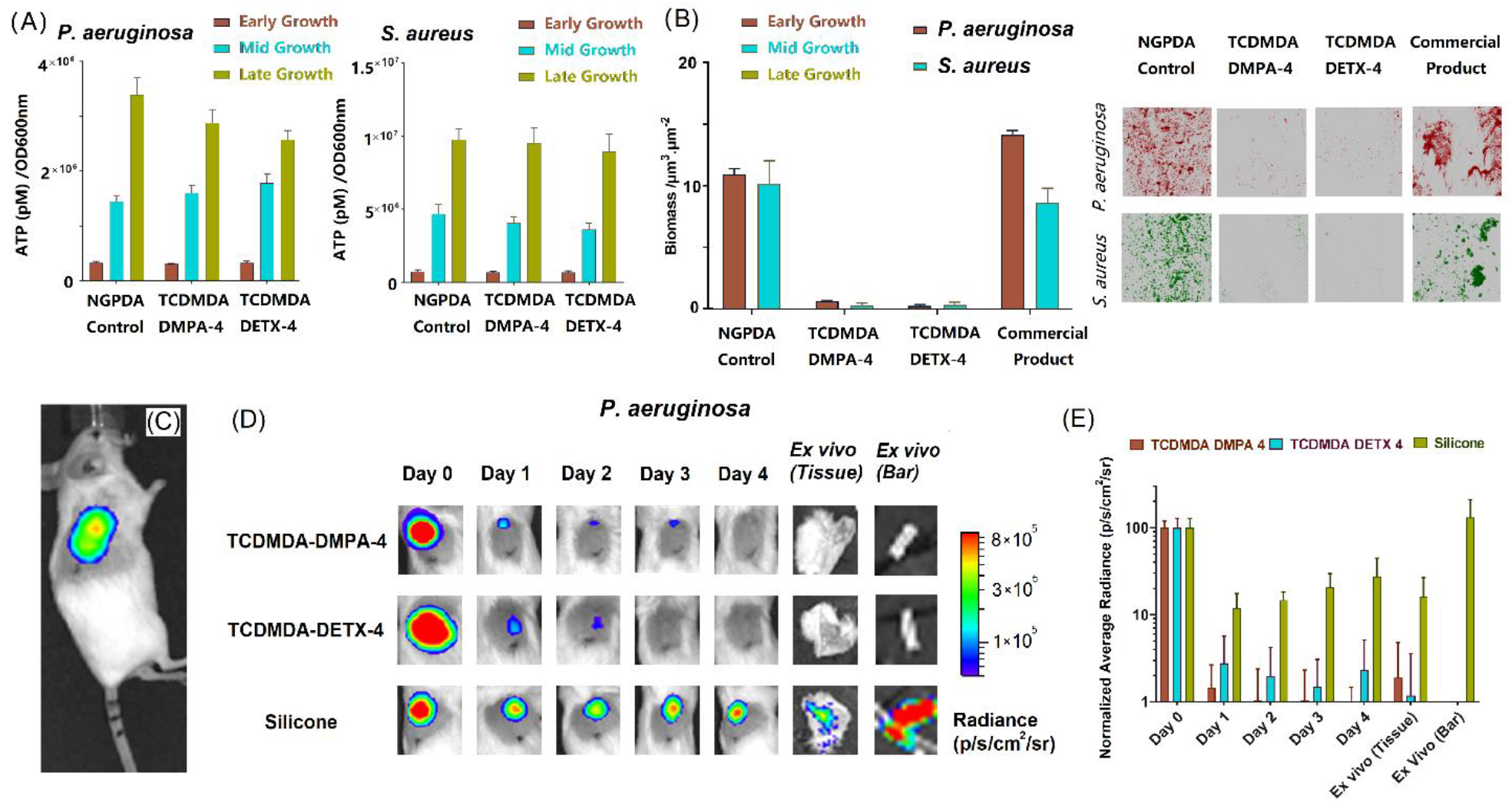
*Assessment of bacterial viability and biofilm formation* in vitro *and infection* in vivo *in a mouse foreign body infection model*. **A)** Bacterial cell viability on printed specimens, RPMI-1460 medium containing the printed sample was inoculated with either *P. aeruginosa* (left) or S. aureus (right) cells. Intracellular ATP levels were quantified at early (OD600nm = 0.25), mid (OD600nm = 0.5) and late (OD600nm = 0.8) exponential phase using a BacTiter-Glo microbial cell viability assay, NGPDA with 4 wt% of DMPA as initiator was used as a control. Data show mean ± standard deviation, n = 3; **B)** Bacterial biofilm formation on printed specimens in vitro: the biofilm biomass of *P. aeruginosa* and *S. aureus* was measured after 72h incubation. Error bars equal ± one standard deviation unit, n = 3. Fluorescent micrographs of mCherry-labelled *P. aeruginosa* (red) and GFP-labelled *S. aureus* (green) growing on each surface (right). mean ± standard deviation, n = 3. Each image is 610 x 610 μm2. **C)** IJ3DP optimized formulations (TCDMDA-DMPA-4 and TCDMDA-DETX-4) and biomedical grade silicone sections (as controls) were implanted subcutaneously in mice. After inoculation, light emission from bioluminescent *P. aeruginosa* at the infection site was measured on the day of inoculation. **D)** Representative bioluminescence outputs overlaid with bright field images of implanted mice infected with *P. aeruginosa* and captured on days 0 to 4. The implanted devices and surrounding tissues were also removed on day 4 from each animal and the device-associated bioluminescence quantified ex vivo. **E)** Bioluminescence was normalised to the output on day 0 showing that the IJ3DP devices were colonized with considerably lower levels of metabolically active bacteria compared with the silicone control.

**Fig. 6:**
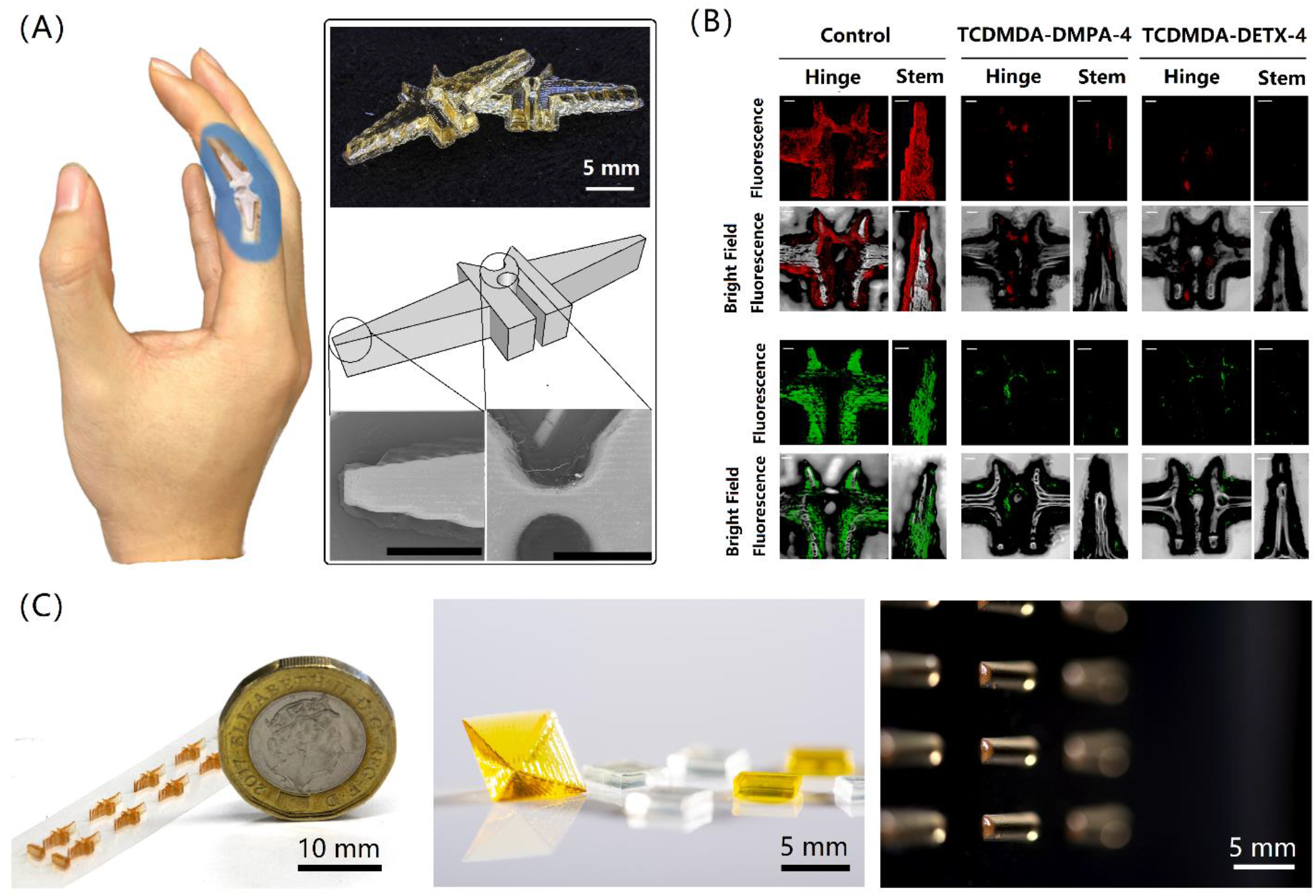
IJ3DP printed finger prosthesis and other demonstrators using the developed ink formulations: **A)** IJ3DP printed finger prosthesis with the developed ink formulations, composed of a central hinge region between two stems, scale bars in the SEM images are 2 mm; **B)** Fluorescence and overlaid fluorescence-brightfield confocal microscopy 3D images showing in vitro biofilm formation imaged using mCherry-labelled *P. aeruginosa* (red) and GFP-labelled *S. aureus* (green) on IJ3DP finger implants with the developed ink formulations. Scale bars represent 200 μm; **C)** specimens of different geometries printed with developed ink using Dimatix printing platform.

### Mechanical Performance Assessment

The two materials selected for scale up were chosen in part due to their likely difference in mechanical properties, allowing a range of moduli to be accessed. The elastic moduli of the printed specimens were determined using dynamic mechanical analysis (DMA), to identify the relationships between the moduli and the composition. Strip specimens with dimensions of 5 x 20 x 1 mm^3^ were printed and tested in tension mode at room temperature (Fig. 2B). The mechanical performance of the printed samples was intimately linked to the photoinitiator concentration, and this effect was pronounced when the DETX initiator was used (usually used for ‘in air’ printing). The variation in photoinitiator concentration led to a modulus for poly-EGDPEA ranging from 0.2 MPa to 180 MPa, while for poly-TCDMDA it ranged between 1.4 GPa to 2.1 GPa. As anticipated, use of poly-TCDMDA and poly-EGDPEA results in a significant separation in the ranges of moduli accessible (Fig. 2B), offering the opportunity to exploit the materials for a range of mechanical requirements such as when integrating with tissues as diverse as cartilage and cancellous bone^[32,33]^.

### Mammalian Cell Cytotoxicity and Attachment

In this study, cytotoxicity assays for mammalian cells (using 3T3 fibroblasts) were conducted using inkjet printed 5 x 5 x 1 mm^3^ cuboid samples (Fig. 3 A and B), following the guidelines presented in ISO 10993 ^[34]^ and detailed in the methods section. These tests were then used as a guide to whether printed constructs could support mammalian cell attachment and proliferation and as a first test of whether these materials could be used safely within the body.

Four sets of samples were considered to have sufficiently low cytotoxicity levels, thus rendering them as appropriate for use. Conditioned media samples from poly-TCDMDA-DETX-4 and poly-TCDMDA-DMPA-4 were the only samples not to exhibit cytotoxicity at any time point, showing lactate dehydrogenase (LDH) levels similar to those of the control (Fig. 3D). Samples from poly-EGDPEA-DMPA-4 and poly-TCDMDA-DMPA-1 showed cytotoxicity over three days, which reduced on subsequent time points. The pattern of photoinitiator content, monomer conversion (discussed in the following section) and cytotoxicity suggests that leaching of residual monomer leads to cytotoxicity, but in the case of EGDPEA-DMPA-4 and poly-TCDMDA-DMPA-1 these are cleared over a timescale of 5 days. All other samples showed either no improvement over the test period, or a highly cytotoxic response indicating that these formulations would be inappropriate for clinical use. Supporting results were obtained using the complementary ‘Presto Blue’ cell viability assay (Fig. S3). Further attachment testing (see methods section) indicated that 3T3 cells attached and proliferated when cultured on poly-EGDPEA-DMPA-4, poly-TCDMDA-DETX-4, poly-TCDMDA-DMPA-1 and poly-TCDMDA-DMPA-4 surfaces (Fig. 3C). Of these, the metabolic activity of cells (determined using the Presto Blue assay) was highest on poly-TCDMDA-DETX-4 and poly-TCDMDA-DMPA-4 (Fig. 3E), closely matching the trends observed for the conditioned cytotoxicity assays.

### Spectroscopic Assessment of Curability

In the above assays, it was hypothesised that the biocompatibility and modulus of the printed material as well as biofilm resistance were influenced by the level of monomer conversion. To investigate this and quantify the relationships, Attenuated Total Reflectance-Infrared Spectroscopy (ATR-IR) was used to determine the residual acrylate content of the films in order to understand whether there was a correlation between photoinitiator concentration, level of conversion and consequently, specimen performance.

Samples were prepared by inkjet printing cuboids of size 5 x 5 x 0.2 mm^3^ thickness. Fig. 4 shows the ATR-IR spectra for polymer samples as a function of the initiator concentration. In the ATR-IR test, the characteristic peak at 810 cm^-1^ (C-H bond out-of-plane bending vibration of the alkene group) indicates the relative amount of unreacted residual alkene group (C=C). For both initiators, the concentration of the residual C=C reduced with increasing initiator concentration, consistent with increased conversion during printing. The relationship between the level of conversion and polymer performance was assessed using the Pearson correlation coefficient (Supplementary Fig.S1). For poly-EGDPEA, the Pearson correlation coefficient between residual alkene groups and mechanical performance, biofilm coverage and cytotoxicity reached 0.82, 0.69 and 0.86 respectively; while for poly-TCDMDA, these values are 0.70, 0.74 and 0.92, thereby confirming the strong link between the residual monomer quantity and key performance measurements.

Combining the correlation analysis for biofilm coverage, mammalian cell cytotoxicity, mechanical performance and level of conversion shows how the performance of printed samples can be optimised by ensuring that all these design criteria are met.

### In Vitro and in Vivo Assessment of the Biofilm Resistance of the Printed Structures

Two ink formulations (poly-TCDMDA-DMPA-4 and poly-TCDMDA-DETX-4) were chosen for further assessment owing to their superior performance across our range of measures. Hemi-cylindrical specimens (7 mm in length and 2 mm in diameter) were printed, matching the dimensions of our control samples, and also enabling sample delivery via a trocar needle in subsequent *in vivo* mouse studies. The viability of *P. aeruginosa* and *S. aureus* in contact with the printed specimens was also tested *in vitro* to ensure that the reduction in biofilm formation was due to colonisation resistance rather than toxicity associated with the material. These experiments revealed no loss of bacterial cell viability (as quantified via intracellular adenosine triphosphate (ATP) levels) during growth in the presence of the candidate samples (Fig. 5A) nor on a printed neopentyl glycol diacrylate (NGPDA) structure that promotes the formation of biofilms as a control polymer ^[20]^. Since no reduction in bacterial viability was observed for bacteria colonizing the positive control, we can conclude that the material itself or any potential residuals in the structure printed by poly-TCDMDA-DMPA-4 and poly-TCDMDA-DETX-4, are not responsible for the lack of biofilm formation. Planktonic bacterial growth experiments (Supplementary Fig.S4) were consistent with the ATP assays and comparable with those in the presence of the NGPDA control.

Quantification of biofilm biomass and the corresponding confocal microscope images are shown in Fig. 5B, which demonstrate the considerable reduction in biofilm biomass observed for both pathogens on the poly-TCDMDA-DMPA-4 and poly-TCDMDA-DETX-4 formulations compared with the poly-NGPDA control device as well as against sample from a commercial silicone rubber finger joint product (OSTF-0, size 0, Osteotec Ltd.) (Fig. 5B).

To understand the printed device’s performance in a more complex host environment, *in vivo* infection experiments were carried out using a murine subcutaneous foreign body implant infection model (Fig.5C). After 4 days of post-surgical recovery, mice were inoculated with a bioluminescent strain of *P. aeruginosa* and the live infected animals imaged daily over 5 days (day 5 to day 9 from implanting; day 0 to day 4 after bacterial inoculation; Fig. 5 D and E), a period over which infection becomes providing *P. aeruginosa* can colonize the implanted device. Light emission from the bioluminescent pathogen demonstrated the presence of metabolically active bacteria at the infection site for all samples at bacterial inoculation day 0 (Fig. 5 D and E). In contrast to the sustained light output indicative of bacterial colonization of the silicone implant, both poly-TCDMDA-DMPA-4 and poly-TCDMDA-DETX-4 formulations showed little bioluminescence (>3 orders of magnitude reduction) consistent with resistance to bacterial attachment *in vivo*. This finding was confirmed by *ex vivo* analysis of the implants and the tissues surrounding the implants after their removal from the mice and re-imaging (Fig. 5 D and E). In contrast to the TCDMDA formulations, the silicone rubber control showed significantly higher bioluminescence consistent with bacterial biofilm formation that also acts as a reservoir for sustaining infection within the interstitial tissues surrounding the implant.

In addition, qualitative imaging of the implants using immunohistochemical staining with antibodies raised against *P. aeruginosa* cells and with the fluorescent dye, FM1-43 (as a marker for host cell and bacterial membranes) revealed evidence of a robust host response and the presence of *P. aeruginosa* cells on both TCDMDA formulations (Supplementary Fig.S5). Given the lack of bioluminescence from such samples, these bacteria are dead, killed via a productive, antibacterial host response since the TCDMDA formulations *per se* are not bactericidal (Fig. 5A and Fig. S5). In contrast, the host defences were unable to kill the *P. aeruginosa* biofilm colonizing the silicone implant given the *in vivo* bioluminescence and *ex vivo* antibody labelling of the bacterial cells (Supplementary Fig. S4). This shows that not only do the devices retain their resistance to bacteria biofilm formation but also drive a productive host response that clears the infecting bacteria.

### Exemplar 3D Printed Biofilm Resistant FingerJoint Prosthetic

To demonstrate that a 3D printed functional device could be realised we chose to manufacture a biofilm resistant finger joint prosthetic using ink jet 3D printing (Fig. 6A) ^[35,36]^. Finger joint prosthetics were printed 1:1 relative to a commercial product using the best performing ink formulations for each monomer (TCDMDA-DMPA-4 and EGDPEA-DMPA-4). The platform used was identical to that used to produce specimens in previous studies, with a typical manufacturing time of around 4 h. The dimension of the printed device was measured from the SEM images and compared with the CAD design (Supplementary Fig. S2). Optimisation of the manufacturing process is needed to ensure such devices could be taken forward for human use, but here we demonstrate that such a route is viable and reliable from the manufacturing aspect.

To check if geometry would impact the functional performance, samples reduced to 1/10 of the original dimensions, were printed and tested *in vitro*. The change in dimension allowed biomass assessment under full view when assessing with fluorescent confocal microscope. The distribution of the *P. aeruginosa* and *S. aureus* biofilms that form on the devices confirmed the cell instructive property was retained (Fig. 6B).

## Discussion

This work has uncovered a series of findings that point towards the reliable manufacturing of cell-instructive structures via IJ3DP. With the advances in high throughput technologies, materials libraries offering diverse functions are becoming widely available ^[21–23, 37]^. It is not trivial, however, to go from an idealized screening approach to materials usable within a manufacturing setting. A key element of the approach outlined has been to establish optimized formulations able to support IJ3DP. In this work, we used a laboratory based single material printing system (Dimatix DMP-2830) with 16 nozzles. On this system, the identified formulations were stable and reliable during printing (Fig. 6C), capable of consistent printing of at least 8 h (the longest print run employed) with no nozzle blockage observed. In the future, this work could be translated to commercial inkjet printing systems that routinely employ multiple printheads containing 1024 nozzles or more, which would result in a throughput at least 14 times greater and with the potential to deploy support structures^[38]^ and therefore more complex geometries. This reliable printing allowed for the manufacture of multiple samples designed for assessment of important characteristics relating to product function. In the case chosen, such a product would need to be safe, meet specified mechanical requirements, and be functional (both structurally and cell instructive). Through a combination of screening to identify candidate formulations and optimization to ensure reliable printing and optimal performance, we were able to direct our activity towards the production of a component that would show high efficacy within the *in vivo* environment. Whilst other longer term host response tests will be required in order to complete the journey towards use in the clinic, our results show that the cell-instructive performance of our selected and optimized formulation are highly promising in an *in vivo* infection model, a critical step towards acceptance. In addition, our choice of manufacturing modality allows for personalization – our experiments utilize a range of shapes and sizes all created with the same platform all supporting our conclusion of material efficacy, and for potential scale up to industrial levels of throughput.

In conclusion, this work demonstrates that advanced cell instructive properties may be incorporated into the production of bespoke medical devices. The materials that we have identified can be employed as feedstock for IJ3DP and that our comprehensive set of *in vitro* and *in vivo* tests confirm the key biofilm resistant property is retained throughout the manufacturing process. Interestingly, our analysis reveals that our selected materials play a previously unobserved role in recruiting host defences that clears the infecting bacteria and prevents biofilm maturation, an exciting finding that deserves future investigation. Whilst the exemplar focuses on the important case of addressing infection while avoiding the opportunity for increasing antimicrobial resistance, our protocol is agnostic with respect to the cell-instructive functionality and may, in principle, be substituted for any other library of materials. This method offers a flexible manufacturing platform suited to the production of medical devices truly tailorable to biological challenge and personalize suitable for translation into clinical practice.

## Materials and Methods

### Ink preparation

All chemicals were purchased from Sigma-Aldrich and used as received. Tricyclo[5.2.1.02,6]decanedimethanol diacrylate (TCDMDA) and Ethylene glycol dicyclopentenyl ether acrylate (EGDPEA) were used as the base monomer in the preparation of all the ink formulations. The photoinitiators used were 2,2-Dimethoxy-2-phenylacetophenone 99% (DMPA) (a type I photoinitiator for nitrogen atmosphere printing) and (2,4-Diethyl-9H-thioxanthen-9-one(DETX), 98%)/(Ethyl 4-(dimethylamino)benzoate(EDB), 99wt%) (a type II photoinitiator system suitable for printing within an air atmosphere). 5 mL of each selected monomer was placed into capped vials (wrapped with aluminum foil) together with a photoinitiator (0.5 wt%, 1 wt%, 2 wt% and 4 wt%) and stirred at 800 rpm at room temperature until the photoinitiator was fully dissolved. The mixture was degassed by purging with nitrogen for 15 min to remove dissolved oxygen. The inks were filtered through a 0.45 μm filter (Minisart, Sartorius Stedim Biotech) in a dark room to remove particulates which may block printer nozzles. In order to maximise printability, inks were sealed and stored at 4°C overnight to help release any bubbles generated during preparation ^[27]^.

### Samples printing

The printing was carried out using a Dimatix DMP-2830. 2 mL of ink was injected into a 10 pL drop volume Dimatix cartridge containing 16 nozzles (21 μm nozzle size). The injection procedure was carried out in a dark room to prevent light-dependent inducing curing. The print cartridge was wrapped in foil to prevent ambient light curing during printing.

Curing was achieved using a UV unit (365 nm and 600 mW/cm2) mounted directly on the printer allowing it to move with the printhead and induce real-time UV illumination and curing contemporaneously with deposition of material.

All the samples with DMPA as a photoinitiator were printed in nitrogen where oxygen levels were controlled to 1 ± 0.5%. The inks with DETX/EDB as initiator were printed in air.

### Polymer mechanical and chemical properties

Dynamic Mechanical Analysis (DMA) tests were carried out at room temperature using a Perkin Elmer DMA 8000 in tension mode. Specimens were printed following a rectangular pattern (20 mm in length and 5 mm in width) with 100 layers. The test length was set to 10 mm and the width and thickness of each sample was measured prior to calculating its modulus. The test period was set to 10 min with 1 Hz extension frequency at room temperature. Infrared Spectroscopy (IR) with an ATR (Perkin Elmer UATR IR) sampling attachment was used to characterize the curability of the printed samples. The spectra for each set of samples were normalized with a peak at 1726 cm^-1^ representing the acrylate carboxyl group. The peak at 810 cm^-1^, which represents the carbon-hydrogen covalent bond on the C=C pairing was used to compare the level of conversion of the printed samples.

### Mammalian Cell Cytotoxicity

Following ISO 10993, Medical Device Tests guidance direct contact (cell attachment test) and indirect extractable testing (cytotoxicity test) was undertaken.

#### Cytotoxicity test

Samples were placed in a 96 well plate, and 1 mL of Industrial Methylated Spirit (IMS, 70% v/v, Fisher Scientific, UK) was added and allowed to evaporate overnight in a microbiological safety cabinet at RT. Samples were washed three times for 5 min each with PBS. Cell culture medium was added (200 μL) to each sample and kept in an incubator at 5% CO2 in air, 37°C. Conditioned medium was collected after 1, 3, 5 and 8 days, and replaced with 200 μL of fresh medium. Cell culture media were prepared by adding 10% (v/v) of Foetal Bovine Serum (FBS, Sigma-Aldrich, UK), 2 mM L-glutamine (Sigma-Aldrich, UK) and 100 U/mL penicillin, 0.1 mg/mL streptomycin and 0.25 μg/mL amphotericin B (Sigma-Aldrich, UK). Immortalized NIH 3T3 mouse embryonic fibroblast cells (3T3s, passage 60) were seeded in a 96 well plate at a density of 5000 cells/well (100 μL) and when they reached confluency, conditioned media were added and cells incubated for a further 24 h at 5% CO2 in air at 37°C. Cells cultured in fresh media were included as a control. The lactate dehydrogenase assay (LDH Assay Kit^®^, Thermo Scientific) and Presto Blue^®^ assay (Invitrogen) were used to test the cytotoxicity of the conditioned media and cell viability, respectively (Supplementary Figure 3). The LDH assay was performed according to the manufacturer’s protocol. Two controls were performed to obtain a spontaneous and a maximum LDH activity. The spontaneous activity was quantified using the medium collected from the controls, where cells were grown in fresh medium. To induce maximum activity, 10 μL of Lysis Buffer (10X) were added to the cells grown in fresh medium for 30 min before assaying.

In brief, 50 μL of each conditioned media sample were transferred to a 96 well plate and 50 μL of the reaction mixture added to each sample, and the plate incubated at room temperature. After 30 min, 50 μL of stop solution was added. The LDH activity was measured by reading the absorbance of the samples at 490 nm (subtracted from the 680 nm reading) using a spectrofluorometer (Tecan Infinite M200 microplate reader). The cytotoxicity of the extracts was calculated using the following equation:

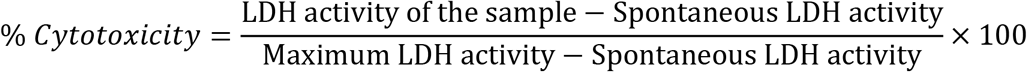

#### Mammalian cell metabolic activity

Presto blue^™^ was diluted 1:10 in cell culture medium and added to the cells. Following aspiration of the medium and washing in PBS for 45 min at 37°C, 5% CO2 in air. The fluorescence intensity of the solution, which is proportional to cellular metabolic activity, was measured at 560 and 590 nm, corresponding to the excitation and emission wavelengths, respectively, and the blank reading (medium without cells) subtracted from each value.

#### Mammalian cell attachment

Samples were placed in a 48 well plate, 1 mL of Industrial Methylated Spirit (IMS, 70% v/v, Fisher Scientific, UK) was added and allowed to evaporate overnight in a microbiological safety cabinet at room temperature. Samples were washed three times for 5 min with PBS. To each sample, 400 μL of cell culture medium was added for 24 h. 3T3 mouse fibroblast cells were seeded on the samples at a concentration of 40,000 cells/well in a total volume of 0.5 mL. After 24 h, the materials were transferred to a new plate to measure the metabolic activity of the cells attached to the scaffold using the Presto Blue assay. Fluorescence intensity was measured and the blank (medium without cells) was subtracted from each value. The test was performed after the cells had been in contact with the test material for 1, 3,5 and 7 days.

#### Live/Dead^®^ cell viability assay

Calcein AM (2.5 μM, representing live cells) and ethidium homodimer-1 (5 μM, red, representing dead cells) were added to the samples and incubated for 30 min at 37°C at 5% CO2, before imaging.

Statistical analyses were performed using Prism 6 (GraphPad Software, v6.01). Two-way ANOVA was performed on cell viability followed by Tukey post-hoc test (n = 3) and on LDH results followed by the Dunnett test (n = 5). A value of p ≤ 0.05 was considered significant. For each condition, mean ± standard deviation was reported.

### Bacterial strains, growth conditions and intracellular ATP assay

*P. aeruginosa* PAO1 (Washington sub-line) and *S. aureus* SH1000 ^[39]^ were routinely grown at 37°C in LB with shaking at 200 rpm or on LB agar (2% w/v). Where required, plasmids for constitutively expressing fluorescent proteins GFP (pBK-miniTn7-egfp) and mCherry (pMMR) were introduced into the relevant host strain by conjugation or electroporation and maintained by supplementing the growth medium with the appropriate antibiotic.

For the quantification of ATP, *P. aeruginosa* and *S. aureus* cell culture samples were taken at early (OD600nm = 0.25), mid (OD600nm = 0.5) or late (OD600nm = 0.8) exponential growth phase. ATP levels were assayed using a BacTiter-GloTM Microbial Cell Viability Assay (Promega UK, Southampton, UK) according to manufacturer’s instructions.

### Bacterial Biofilm Formation

Bacterial biofilm formation assays were conducted as follows. Briefly, UV-sterilized IJ3DP devices (cuboids, tablets or finger implants) with the developed formulations were incubated with bacteria at 37°C with 60 rpm shaking for 72h in RPMI-1640. Air-dried samples were examined using a Carl Zeiss LSM 700 laser scanning confocal microscope fitted with 405 nm, 488 nm and 555 nm excitation lasers and a 10x/NA 0.3 objective. Images were acquired using ZEN 2009 imaging software (Carl Zeiss). Bacterial surface coverage was quantified using Image J 1.44 software (National Institutes of Health, USA) and Comstat B ^[40]^.

### Mouse foreign body infection model

All animal experiments were approved following local ethical review at the University of Nottingham and performed under Home Office licence 30/3082. Female BALB/c mice, 19-22g (Charles River; 3 mice per infected implant and 2 mice per uninfected implant control) were housed in individually vented cages under a 12 h light cycle, with food and water *ad libitum*. *P. aeruginosa* (strain PAO1-L CTX:*lac-lux*) was grown overnight in LB broth at 37°C, diluted 1:100 in LB and grown at 37°C to mid-log phase (OD600). The cultures were washed in PBS+10 % v/v glycerol and aliquots stored at - 80°C. When required, aliquots were removed, serially diluted and cultured on LB agar plates and the number of colony forming units (CFUs) determined. One hour before device implantation via a Trocar needle, Carprofen (2.5mg/kg) was administered by subcutaneous injection to reduce pain and inflammation. Animals were anaesthetised with 2% isoflurane, their flanks shaved and the skin cleaned with Hydrex surgical scrub. A small incision was made and the catheter implanted via a 9g trocar needle and closed with Gluture skin glue (Abbott Laboratories). Mice were allowed to recover for 4 days. Under anaesthesia, 10^5^ bioluminescent *P. aeruginosa* cells in 20 μl PBS were injected into the IJ3DP devices implanted in the mice. The progress of bacterial infection was imaged as bioluminescence using an IVIS^™^ Spectrum (Perkin Elmer). The infected animals were tracked daily for 5 days via whole animal imaging for the presence of metabolically active bacteria at the infection site. After sacrificing the mice, the IJ3DP devices and the surrounding tissues were removed and reimaged *ex vivo* using an IVIS^™^ Spectrum to quantify bacterial bioluminescence. In addition, the implants were fixed with 10% formal saline and subjected to immunohistochemical analysis and confocal microscopy using antibodies raised against *P. aeruginosa* cells (Invitrogen) and detected using a secondary goat anti-mouse fluorescent conjugate (quantum dot 705; Thermofisher). Total host cell and bacterial membrane biomass on the implants was stained using the fluorescent cell membrane probe, FM1-43 (Thermofisher). After staining implants were imaged using by confocal fluorescence microscopy (Zeiss LSM700).

## Acknowledgments

YH, RW, CT, RH, and BB were funded by Engineering and Physical Sciences Research Council grants EP/I033335/2, EP/N024818/1 and EP/P031684/1, EP and FR by EP/L015072/1 and MA, PW, J-FD by the Wellcome Trust Senior Investigator Joint Awards to 103882/Z/14/Z and 103884/Z/14/Z. Funding for open access charge was provided by UK Research and Innovation.

The manuscript was written with contributions of all authors. The majority of the experimental work was carried out by YH. Support for 3D printing was provided by BB, RW, CT and RH. JL conducted *in vivo* mouse experiments and J-FD and AH conducted the *in vitro* bacterial assays overseen by PW. Cytotoxicity experiments were conducted by EP under the supervision of FR. Chemical characterisation and materials understanding was overseen by DI. The work was conceived and organised by RW, MA and PW.

## Supplementary Materials

**Table S1:**
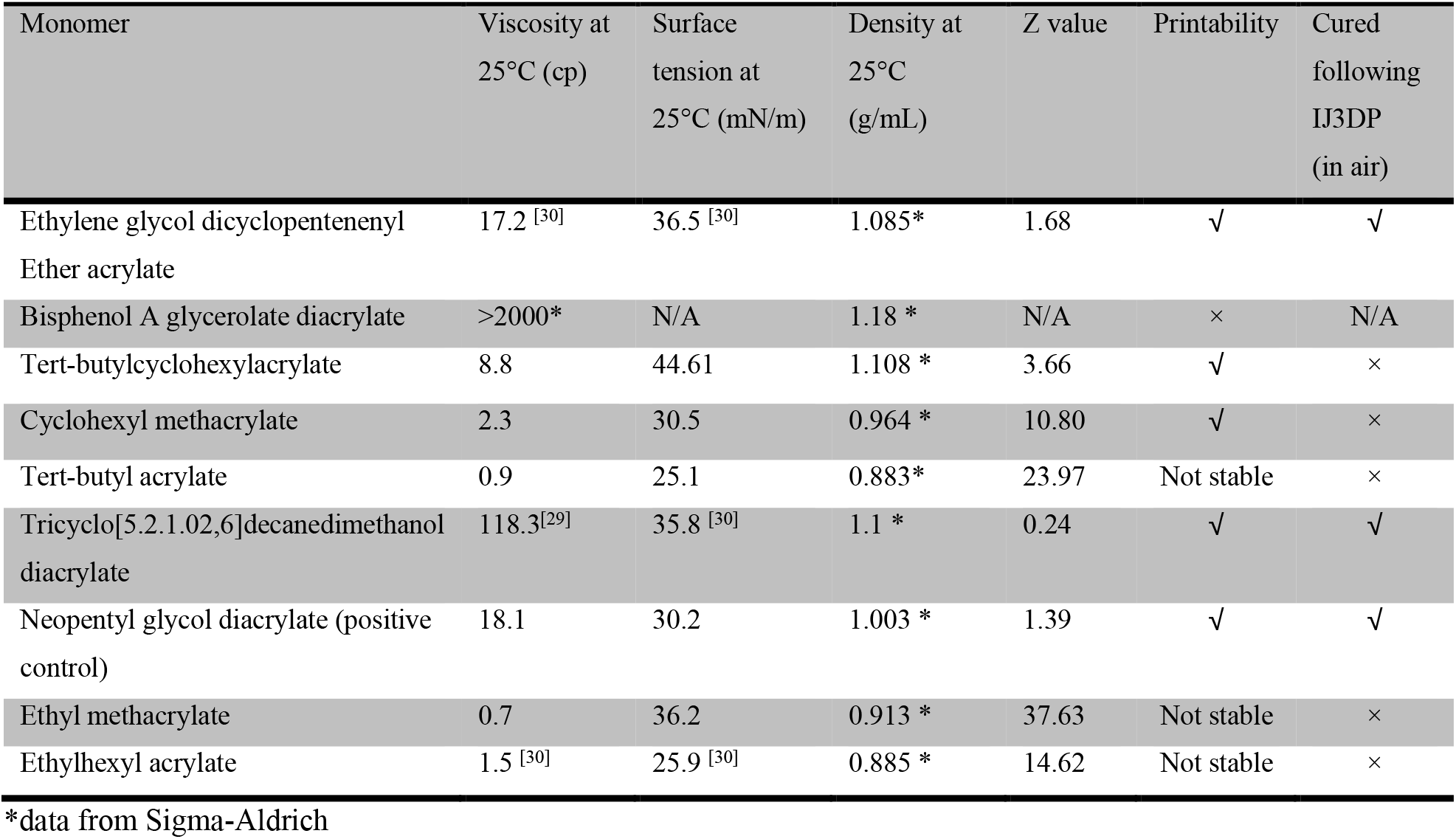
Monomers that have shown resistance to bacterial biofilm resistance after polymerization were selected from polymer library and tested using our Dimatix DMP 2830 printing platform to assess if they are printable and curable in IJ3DP process. Printability is assumed within the range 1 < Z < 10 ^[27]^ and then an assessment of whether reliable and consistent printing was possible was made by manual observation of the droplet formation and deposition. Curing was assessed by observing whether a 3D printed material was self-supporting.

| Monomer | Viscosity at 25°C (cp) | Surface tension at 25°C (mN/m) | Density at 25°C (g/mL) | Z value | Printability | Cured following IJ3DP (in air) |
| --- | --- | --- | --- | --- | --- | --- |
| Ethylene glycol dicyclopentenyl Ether acrylate | 17.2 [30] | 36.5 [30] | 1.085* | 1.68 | ✓ | ✓ |
| Bisphenol A glycerolate diacrylate | >2000* | N/A | 1.18 * | N/A | × | N/A |
| Tert-butylcyclohexylacrylate | 8.8 | 44.61 | 1.108 * | 3.66 | ✓ | × |
| Cyclohexyl methacrylate | 2.3 | 30.5 | 0.964 * | 10.80 | ✓ | × |
| Tert-butyl acrylate | 0.9 | 25.1 | 0.883* | 23.97 | Not stable | × |
| Tricyclo[5.2.1.02,6]decanedimethanol diacrylate | 118.3[29] | 35.8 [30] | 1.1 * | 0.24 | ✓ | ✓ |
| Neopentyl glycol diacrylate (positive control) | 18.1 | 30.2 | 1.003 * | 1.39 | ✓ | ✓ |
| Ethyl methacrylate | 0.7 | 36.2 | 0.913 * | 37.63 | Not stable | × |
| Ethylhexyl acrylate | 1.5 [30] | 25.9 [30] | 0.885 * | 14.62 | Not stable | × |

\*data from Sigma-Aldrich

**Table S2:** The composition of ink formulations and their abbreviations: Tricyclo[5.2.1.02,6]decanedimethanol diacrylate (TCDMDA), Ethylene glycol dicyclopentenyl ether acrylate (EGDPEA), 2,2-Dimethoxy-2-phenylacetophenone (DMPA), (2,4-Diethyl-9H-thioxanthen-9-one(DETX), Ethyl 4-(dimethylamino)benzoate(EDB), neopentyl glycol diacrylate (NGPDA)

| Acronym | Monomers |  |  | Initiators |  |  |
| --- | --- | --- | --- | --- | --- | --- |
|  | TCDMDA | EGDPEA | NGPDA | DETX | EDB | DMPA |
| TCDMDA-DETX-0.5 | √ | -- | -- | 0.5wt% | 0.5wt% | -- |
| TCDMDA-DETX-1 | √ | -- | -- | 1wt% | 1wt% | -- |
| TCDMDA-DETX-2 | √ | -- | -- | 2wt% | 2wt% | -- |
| TCDMDA-DETX-4 | √ | -- | -- | 4wt% | 4wt% | -- |
| TCDMDA-DMPA-0.5 | √ | -- | -- | -- | -- | 0.5wt% |
| TCDMDA-DMPA-1 | √ | -- | -- | -- | -- | 1wt% |
| TCDMDA-DMPA-2 | √ | -- | -- | -- | -- | 2wt% |
| TCDMDA-DMPA-4 | √ | -- | -- | -- | -- | 4wt% |
| EGDPEA-DETX-0.5 | -- | √ | -- | 0.5wt% | 0.5wt% | -- |
| EGDPEA-DETX-1 | -- | √ | -- | 1wt% | 1wt% | -- |
| EGDPEA-DETX-2 | -- | √ | -- | 2wt% | 2wt% | -- |
| EGDPEA-DETX-4 | -- | √ | -- | 4wt% | 4wt% | -- |
| EGDPEA-DMPA-0.5 | -- | √ | -- | -- | -- | 0.5wt% |
| EGDPEA-DMPA-1 | -- | √ | -- | -- | -- | 1wt% |
| EGDPEA-DMPA-2 | -- | √ | -- | -- | -- | 2wt% |
| EGDPEA-DMPA-4 | -- | √ | -- | -- | -- | 4wt% |
| NGPDA-DMPA-0.5 | -- | -- | √ | -- | -- | 0.5wt% |
| NGPDA-DMPA-1 | -- | -- | √ | -- | -- | 1wt% |
| NGPDA-DMPA-2 | -- | -- | √ | -- | -- | 2wt% |
| NGPDA-DMPA-4 | -- | -- | √ | -- | -- | 4wt% |

### Correlation Analysis

Statistical analysis was performed using GraphPad Prism 6: Pearson’s correlation coefficient was introduced to quantify the correlation of the residual C=C groups versus bacterial surface coverage, mammalian cell cytotoxicity and storage modulus of the printed polymeric structure.

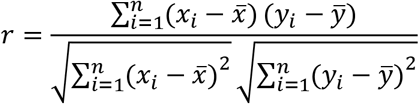

where *n* is the sample size, *x_i_* and *y_i_* are the single data points, 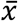 and *ȳ* are the mean value.

**Fig. S1:**
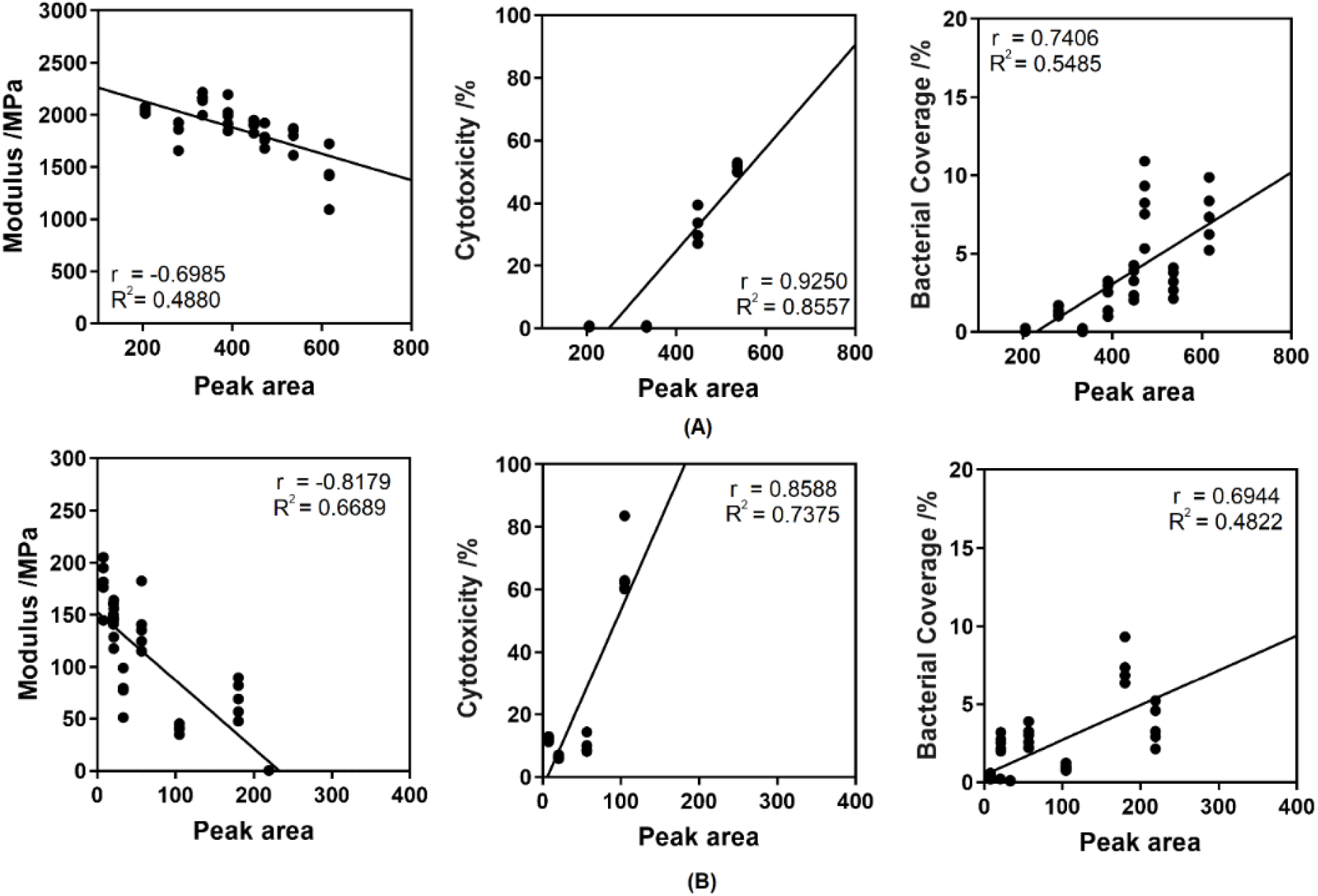
Pearson’s correlation analysis between the C=C residual level and bacterial biofilm formation, cytotoxicity and storage modulus respectively. The residual monomer level was judged by the peak area of C-H out-of-plate bending vibration on C=C at 810 cm^-1^. (A) TCDMDA-DMPA. (B) EGDPEA-DMPA

### Comparison of Specimens dimensions with CAD design

**Fig. S2:**
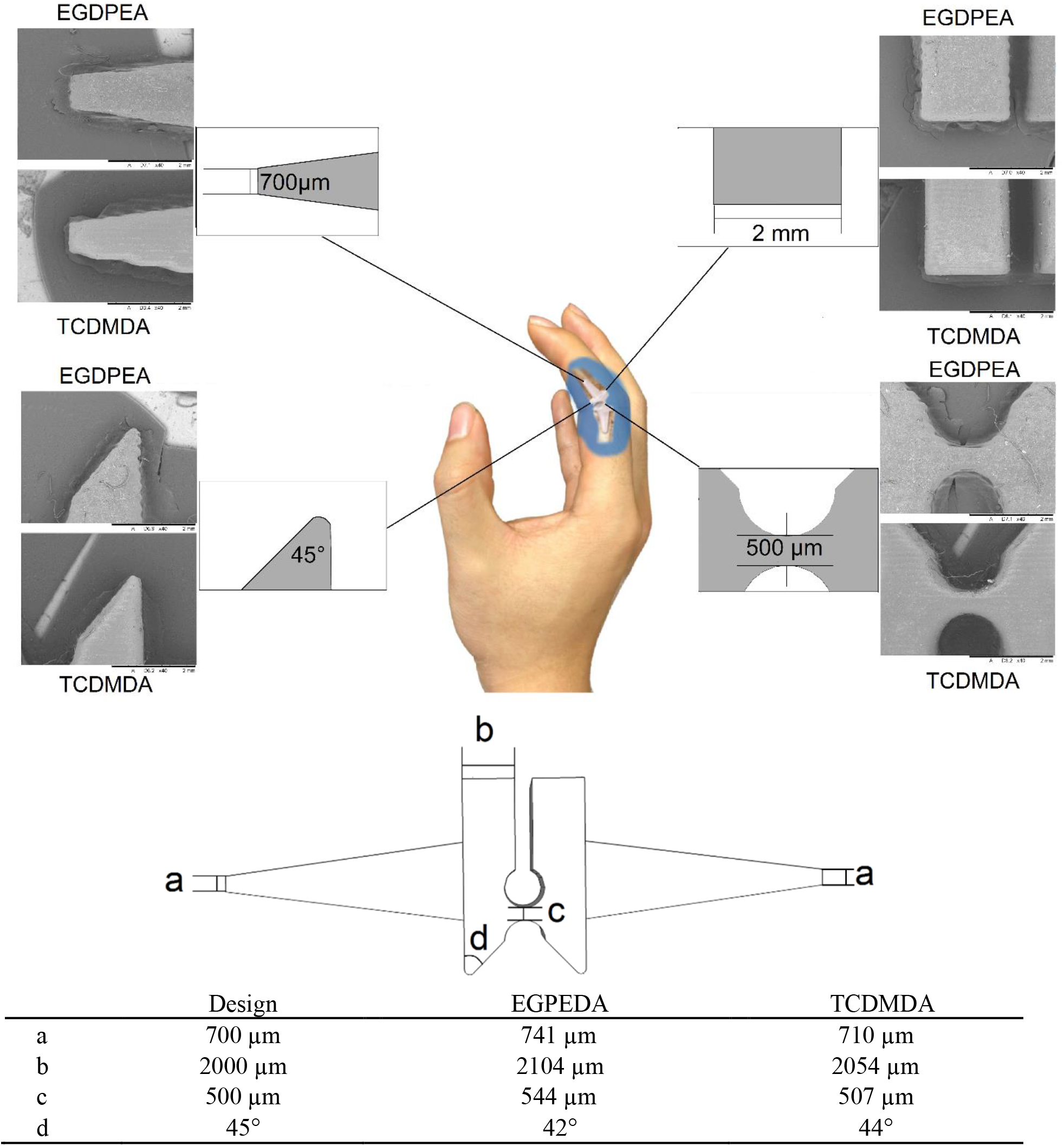
Comparision of the CAD designed feature size with the actual printed specimen size from 4 different key feature points (a-d).

### Mammalian cell cytotoxicity

**Fig. S3:**
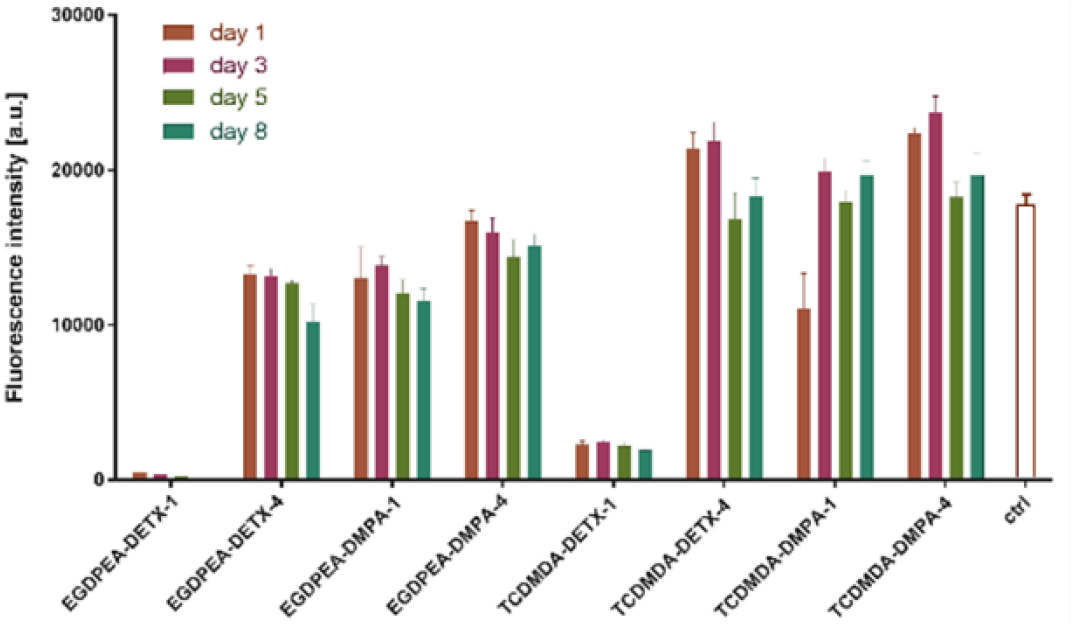
Mammalian cytotoxicity assays using the Presto Blue^®^ assay for printed TCDMDA and EGDPEA samples with both DMPA and DETX initiators at 1 wt % and 4 wt %. The sampling time was 1, 3, 5 and 8 days. The data are presented are mean ± standard deviation, n=5 (*p ≤ 0.05).

### Bacterial Viability test

**Fig. S4:**
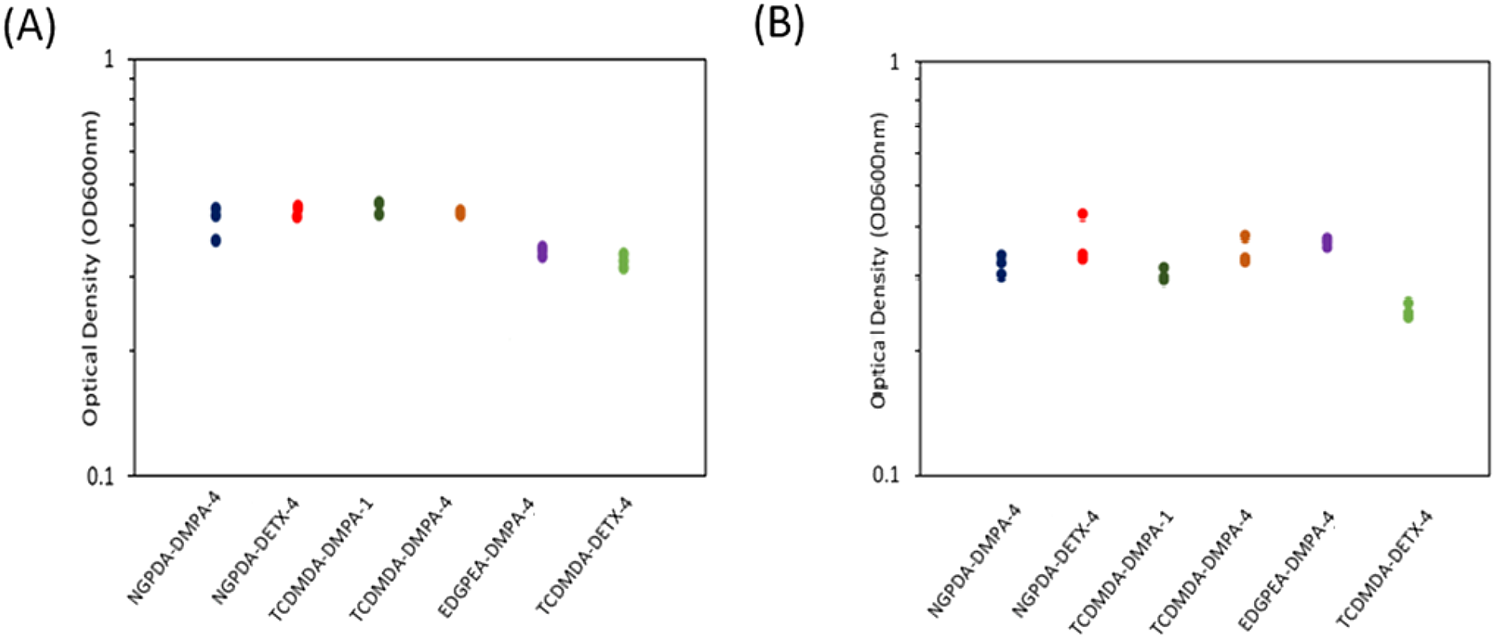
Devices were printed with different ink formulations: NGPDA-DMPA-4, NGPDA-DETX-4, EDGPEA-DMPA-4, TCDMDA-DMPA-1, TCDMDA-DMPA-4, or TCDMDA-DETX-4. Samples were immersed in RPMI-1460 medium inoculated with *P. aeruginosa* (**A**) or *S. aureus* (**B**) cells for 24 h. The stationary phase OD_600_ reached for *P. aeruginosa* and *S. aureus* are shown in (A) and (**B**) respectively. Mean ± Standard Deviation, n = 3.

### Bacterial and antibody marking of implant tested in vivo

**Fig. S5.**
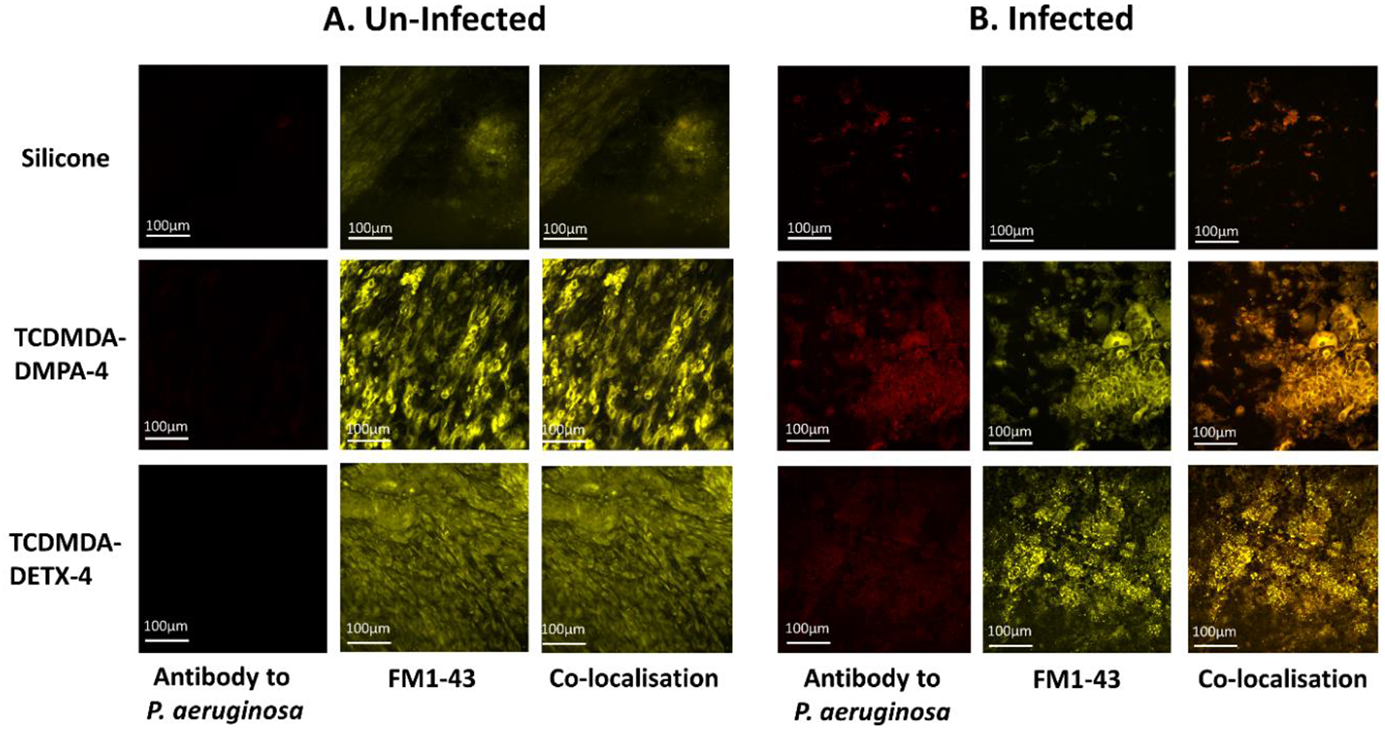
Immunohistochemistry of the TCDMDA and silicone implants *ex vivo* showing the presence of bacteria and host cells. TCDMDA and silicone implants were recovered from (A) control, uninfected mice and (B) mice infected with *P. aeruginosa*. Implants were stained with an antibody to *P. aeruginosa (red)* and with the membrane stain FM1-43 (yellow). No bacteria were detected on the uninfected controls. On silicone, large numbers of individual whole bacterial cells and some bacterial aggregates are apparent. For both TCDMDA formulations but not silicone, there is a strong host response that co-localizes with bacterial cells and cell fragments.

## Notes

### Competing Interest Statement

The authors have declared no competing interest.

